# Analysis of biosynthesis and composition of cuticular wax in wild type bilberry (*Vaccinium myrtillus* L.) and its glossy mutant

**DOI:** 10.1101/2020.04.01.019893

**Authors:** Priyanka Trivedi, Nga Nguyen, Linards Klavins, Jorens Kviesis, Esa Heinonen, Janne Remes, Soile Jokipii-Lukkari, Maris Klavins, Katja Karppinen, Laura Jaakola, Hely Häggman

## Abstract

Cuticular wax plays an important role in fruits in protection against environmental stresses and desiccation. In this study, biosynthesis and chemical composition of cuticular wax in wild type (WT) bilberry fruit was studied during development and compared with its natural glossy type (GT) mutant. The cuticular wax load in GT fruit was comparable to WT fruit. In both fruits, triterpenoids were the dominant wax compounds with decreasing proportion during the fruit development accompanied with increasing proportion of aliphatic compounds. Gene expression studies supported the pattern of compound accumulation during fruit development. Genes *CER26-like, FAR2, CER3-like, LTP, MIXTA*, and *BAS* exhibited prevalent expression in fruit skin indicating role in cuticular wax biosynthesis and secretion. In GT fruit, higher proportion of triterpenoids in cuticular wax was accompanied by lower proportion of fatty acids and ketones compared to WT fruit as well as lower density of crystalloid structures on berry surface. Our results suggest that a marked reduction in ketones in cuticular wax may play a significant role in the formation of glossy phenotype leading to the loss of rod-like structures in epicuticular wax layer of GT fruit.

**Highlight:** Chemical composition and morphology of cuticular wax along with gene expression for wax biosynthetic genes varied between glossy type mutant (GT) and wild type (WT) fruit.

## Introduction

Cuticle is a lipophilic layer on aerial parts of plant surface, composed of cuticular wax and cutin, a polyester polymer matrix. Cuticle plays an important role in preventing water loss, protection against UV radiation and pathogen attack in plants, including fruits at different developmental stages and during storage period (Lara *et al*., 2014; Petit *et al.*, 2017). Cuticular wax is a complex mixture of very long chain fatty acids (VLCFAs) and their derivatives, such as aldehydes, alkanes, ketones, primary and secondary alcohols, esters as well as secondary metabolites, including triterpenoids, sterols, and phenolic compounds (Kunst and Samuels, 2009; Lara *et al*., 2015). Fruit cuticular waxes have especially been shown as good sources of triterpenoids, which are well known for their health beneficial properties, including antioxidant and anti-inflammatory properties as well as decreasing risk for cardiovascular diseases (Szakiel *et al*., 2012; Han and Bakovic, 2015).Previous studies have shown that the composition of cuticular wax varies not only between species, cultivars and organs, but also with the developmental stage of the same organ (van Maarseveen *et al*., 2009). A variable trend in wax deposition rate as well as alterations in chemical composition of cuticular wax through fruit development in various species have been reported (Curry, 2005; Domínguez *et al*., 2008; Wang *et al*., 2016; Trivedi *et al*., 2019b).

Cuticular wax can be seen as whitish (glaucous) or glossy epicuticular wax, while it is also embedded on the cutin as intracuticular wax (Jenks *et al*., 2002; Ensikat *et al*., 2006). The chemical basis for the difference between glaucous and glossy wax phenotypes is unclear although has been studied in various species. Glaucous leaf and stem mutants of Arabidopsis showed higher wax load accompanied by higher density of epicuticular wax crystals (Jenks *et al* 1996). Characterization of naturally occurring glaucous lines have identified β-diketones to be responsible for glaucousness in wheat and barley (Hen-Avivi *et al*., 2016). Among fruits, orange glossy type mutant fruits showed a decrease in wax load accompanied by reduction in proportion of aldehydes affecting crystalloid formation (Liu *et al*., 2012; 2015). In cucumber, *CsCER1*-RNAi transgenic lines showing glossy phenotype demonstrated inhibited wax crystallization attributed to decrease in proportion of alkanes as compared to wild type lines (Wang *et al*., 2015b). In case of apples, glossiness (or greasiness) was attributed to melting of wax crystalloids and formation of amorphous wax (Yang *et al*., 2017). There is a need of more fruit specific studies to understand the chemical and morphological basis of glossy and glaucous phenotypes.

The wax biosynthesis pathways with key genes have been elucidated by studies performed especially in Arabidopsis. In general, the biosynthesis of aliphatic compounds of cuticular wax starts from *de novo* fatty acid biosynthesis in plastids producing C_16_–C_18_ fatty acids by β-ketoacyl-ACP synthase (KAS) as key enzyme (Fig. S1). The later stages of biosynthesis occur in endoplasmic reticulum (ER) exclusively in epidermal cells where elongation of VLCFAs (C_20_–C_34_) is facilitated by β–ketoacyl-CoA-synthase (KCS). The different classes of aliphatic compounds of the cuticular wax are modified from the VLCFAs by two pathways; acyl reduction pathway (alcohol forming) to produce primary alcohols and wax esters, and decarbonylation pathway (alkane forming) to produce aldehydes, alkanes, ketones, and secondary alcohols. The primary alcohols are biosynthesized by fatty acyl-CoA reductase (FAR) encoded by *CER4* (Rowland *et al*., 2006), and then further esterified to wax esters by wax synthase enzyme (WSD1/DGAT). *CER1* and *CER3*, encoding aldehyde decarbonylase and VLC-acyl-CoA reductase, respectively, have been identified to be involved in alkane synthesis (Rowland *et al*., 2007; Bernard *et al*., 2012). Secondary alcohols are produced from alkanes by mid-chain alkane hydrolase (MAH1). The wax components are transported to Golgi (McFarlane *et al*., 2014) and exported through the plasma membrane by heterodimer ABCG transporter family proteins, known as ABC11/WBC11 and ABC12/CER5 in Arabidopsis (Bird *et al*., 2007). The wax compounds are transported and secreted to the cell wall by non-specific lipid transfer protein (LTP; Kunst and Samuels, 2009). However, the mechanism of wax secretion is not yet fully understood. The wax triterpenoids are biosynthesized from squalene and cyclized by oxidosqualene cyclase enzymes (OSCs) such as β-amyrin synthase (BAS) and lupeol synthase (LUS), to produce variety triterpenoids and steroids (Fig. S1; Delis *et al*., 2011).

There are only few studies of wax biosynthesis in fruits and the studies have mostly focused on horticultural plants, such as tomato (*Solanum lycopersicum* L., Mintz-Oron *et al*., 2008), sweet cherry (*Prunus avium* L., Alkio *et al*., 2012), apple (*Malus domestica* L., Albert *et al*., 2013), orange (*Citrus sinensis* L., Liu *et al*., 2015; Wang *et al*., 2016), mango (*Mangifera indica* L., Tafolla-Arellano *et al*., 2017), and cucumber (*Cucumis sativus* L., Wang *et al*., 2015a,b; Wang *et al*., 2018). Bilberries (*Vaccinium myrtillus* L.) are deciduous shrubs with wide distribution in cool temperate regions and mountain areas of Europe and Asia. As an abundant resource in Northern forest, wild bilberries play a significant role in food industry. The berries provide also an excellent raw material for extraction of health beneficial products, like anthocyanins, but the leftovers of food industry (berry press cakes) can also be utilized for extraction of bioactive wax compounds (Lara *et al*., 2014; Trivedi *et al*., 2019a).

The goal of this study was to explore wild type bilberry fruit (WT) and glossy type natural mutant (GT) for differences in composition, morphology and biosynthesis of cuticular wax through developmental stages. We studied overall wax amounts, proportion of wax compound classes and absolute wax amounts (in µg/cm^2^) in WT and GT through developmental stages. To put compositional data into context, we identified genes related to cuticular wax from *de novo* bilberry transcriptome constructed earlier (Nguyen *et al*., 2018) and used as an exploratory data to understand the wax biosynthesis in bilberry.

## Materials and methods

### Plant materials

Wild type (WT) and glossy type mutant (GT) fruits of bilberry (*Vaccinium myrtillus* L.) at four developmental stages, named S2 (small green fruits), S3 (large green fruits), S4 (ripening red fruits), and S5 (fully ripe blue fruits), as described previously (Nguyen *et al*., 2018), were utilized for studies (Fig. 1). The fruits were collected using forceps during June to August 2018 from the natural forest stand in Oulu, Finland (65°03’37.0”N 25°28’30.4”E).

**Fig. 1.**
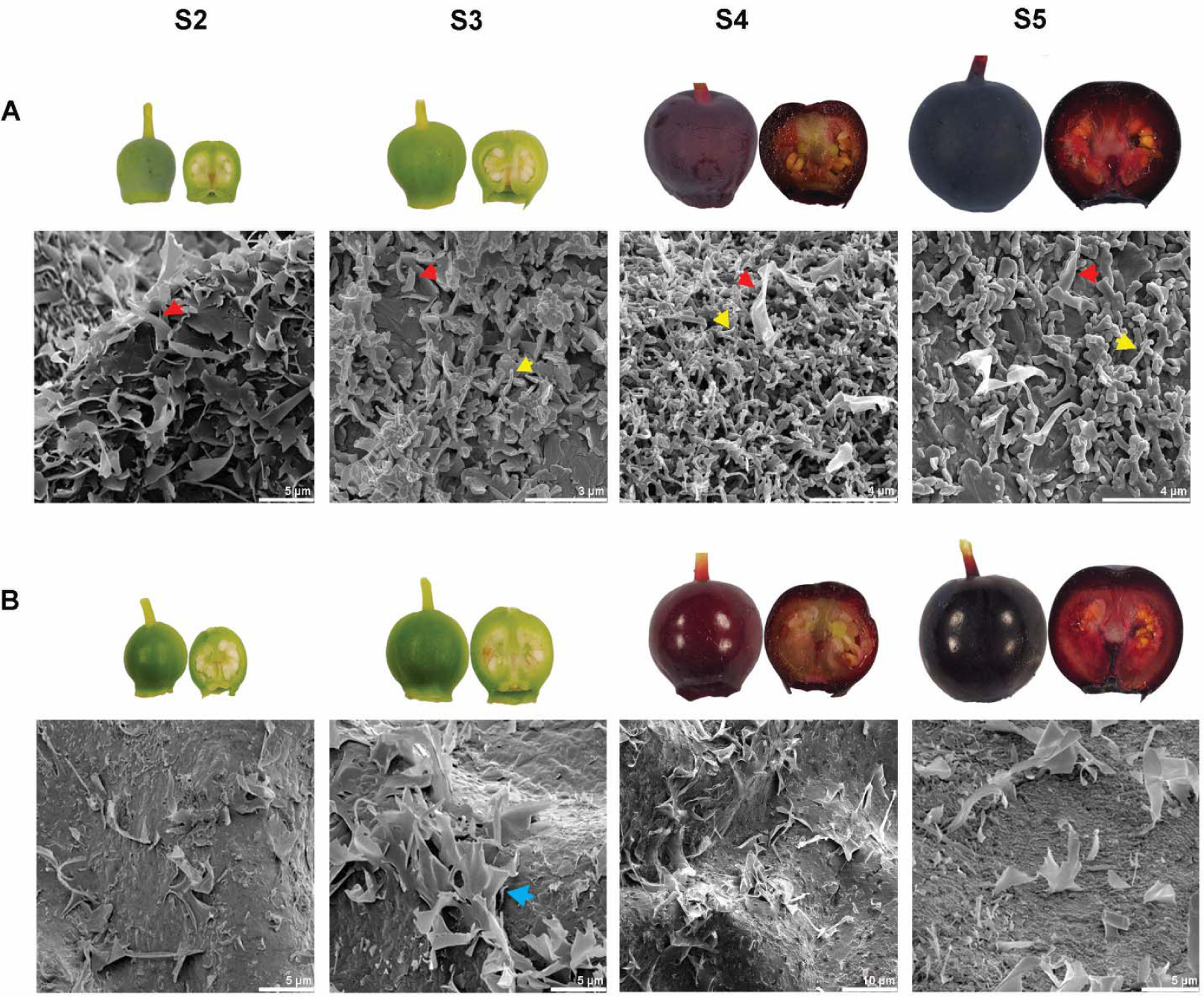
Changes in epicuticular wax morphology on the surface of (A) wild type (WT) and (B) glossy type mutant (GT) bilberry fruits during development. Red arrows indicate platelet structure, yellow arrows indicate rod-like structure, blue arrows indicate membranous platelet structure. S2, small green fruits; S3, large green fruits; S4, ripening red fruits; S5, fully ripe blue fruits.

### Scanning electron microscopy (SEM)

For SEM analysis, fresh berries were dried immediately after collection by using a vacuum freeze-drier (Edwards High Vacuum International, West Sussex, England) before fixed on aluminium stubs. The berry surfaces were sputter-coated with 20 nm layer of platinum by using a sputter coater (Agar High Resolution Sputter Coater, Agar Scientific Ltd, Essex, UK) and then investigated for the three-dimensional surface micromorphology by using SEM (Helios Nanolab 600, Oregon, USA).

### Cuticular wax extraction and determination of wax amount

Immediately after collection, the cuticular wax from the four developmental stages of both WT and GT fruits was separately extracted with chloroform (Sigma-Aldrich, St. Louis, USA). Berries were dipped in 15 mL chloroform for 1 min. The extract was evaporated to dryness under nitrogen flow at room temperature followed by the measurement of dry weight. The cuticular wax extraction was performed in triplicates for each berry developmental stage (except glossy type mutant S4 stage, where due to unavailability of glossy type mutants, extraction was performed in duplicates). The amount of wax was expressed as weight per unit surface area (µg/cm^2^). For calculating the surface areas, images of the dipped berries on a white surface were taken immediately after wax extraction. Image J software v1.50i (NIH, Maryland, USA) was used to calculate the total surface area of the berries as S= 4 πr^2^, where r is the radius of berry (assuming that the berries are spherical).

### GC-MS analysis

Derivatization of fatty acids and GC-MS analysis was performed as described previously by Trivedi *et al.* (2019a). GC-MS analysis was performed using a PerkinElmer Clarus 580 system equipped with a Clarus SQ 8 C mass-selective detector (Waltham, MA, USA) and an Omegawax 250 column (30 m × 0.25 mm, 0.25 µm, Darmstadt, Germany). Analysis of FAME’s and polyfunctional compounds as trimethylsilyl derivatives was performed on an Elite-5MS column (30 m × 0.25 mm, 0.25 μm, PerkinElmer). Identification of compounds was done using NIST MS 2.2 library (Gaithersburg, MD, USA). The analysis was performed in triplicate.

### Identification of candidate genes related to the wax biosynthesis

*De novo* transcriptome database of bilberry (Nguyen et al., 2018), was utilized for identifying candidate genes related to wax biosynthetic pathway. The identity of the genes were verified by BLASTX with threshold E-value cut off of 1e-5 against reference protein sequences of Arabidopsis (The Arabidopsis Information Resource - TAIR, https://www.arabidopsis.org/) and other fruits (National Centre for Biotechnology Information - NCBI).

### RNA extraction and qRT-PCR

Skin and pulp were separated from the four developmental stages of both WT and GT fruits by using a razor blade. After sectioning, the pulp and skin samples were immediately frozen in liquid nitrogen and stored at -80 °C until used for RNA extraction. For RNA extraction, tissues were ground to fine powder under liquid nitrogen. Total RNA was extracted with three biological replicates following the protocol of Jaakola *et al*. (2001). The quantity and quality of RNA samples were tested by Nanodrop (Thermo Scientific) and 1% agarose gel stained with ethidium bromide. Then, cDNA was synthesized from 5 µg of total RNA using Superscript III Reverse Transcriptase (Invitrogen,Carlsbad, CA, USA) according to the manufacturer’s instructions. The cDNA was purified from genomic DNA as described by Jaakola *et al.* (2004).

The qRT-PCR analysis was performed with LightCycler 480 instrument and software v1.5.0.39 (Roche Applied Sciences, Foster, CA, USA). The transcript abundance was detected by using LightCycler® SYBR Green I Master qPCR kit (Roche). The qRT-PCR conditions were 95□°C for 10□min followed by 45 cycles of 95□°C for 10□s, 60□°C for 10□s and 72□°C for 20□s. The qRT-PCR results were calculated by LightCycler® 480 software (Roche), using the calibrator-normalized PCR efficiency-corrected method (Technical note no. LC 13/2001, Roche). Glyceraldehyde-3-phosphate dehydrogenase gene (*GAPDH*, GenBank accession number AY123769) was used as internal control to normalize the relative transcript levels. The expression of *GAPDH* has been shown to be stable during the bilberry fruit development (Jaakola *et al.*, 2002). Gene-specific primer sequences used for qRT-PCR analysis are listed in Table S1.

### Statistical analysis

Significant differences in various compound classes between WT and GT fruit at p-value < 5% were analyzed by independent sample *t*-test using SPSS Statistic program v26. The relative means of expression of the studied genes in WT and GT fruit were compared with either *t*-test or Mann-Whitney U test using R v3.6.2 (R Core Team, 2019).

## Results

### Cuticular wax morphology

By visual inspection of fruit surface, the difference in appearance between glaucous WT and GT bilberry can be detected already in early stage (S2) of fruit development (Fig. 1). SEM analysis of fruit surface during WT fruit development showed a dense cover of irregular platelets at S2 stage (Fig. 1A). At S3, S4 and S5 stages of WT fruit development, a syntopism of dense rod-like structures with irregular platelets was seen. In the GT fruit, an amorphous layer of wax with markedly lower density of crystalloid structures compared to WT bilberry fruit was detected throughout the fruit development (Fig. 1B). Only membranous platelets but no rod-like structures were detected in GT fruit.

### Cuticular wax load

Both WT and GT bilberry fruit had cuticular wax present already in S2 stage (Fig. 2). The amount of wax per berry was found to gradually increase during fruit development of both WT and GT fruit reaching in ripe stage (S5) the amount of 367.6 μg in WT fruit and 315.5 μg in GT fruit (Fig. 2A). No marked differences in the total wax amount between the WT and GT fruits in any developmental stage was detected. Wax amount per surface area increased slightly in both WT and GT fruit at ripening stage (S4) while slight decrease towards S3 and S5 stages was detected (Fig. 2B). The measured surface areas of GT fruits at S4 and S5 stages were slightly smaller than WT berries explaining the somewhat higher wax amount per berry in WT berries in S4 and S5 stages that could not be seen when wax amount was expressed per surface area.

**Fig. 2.**
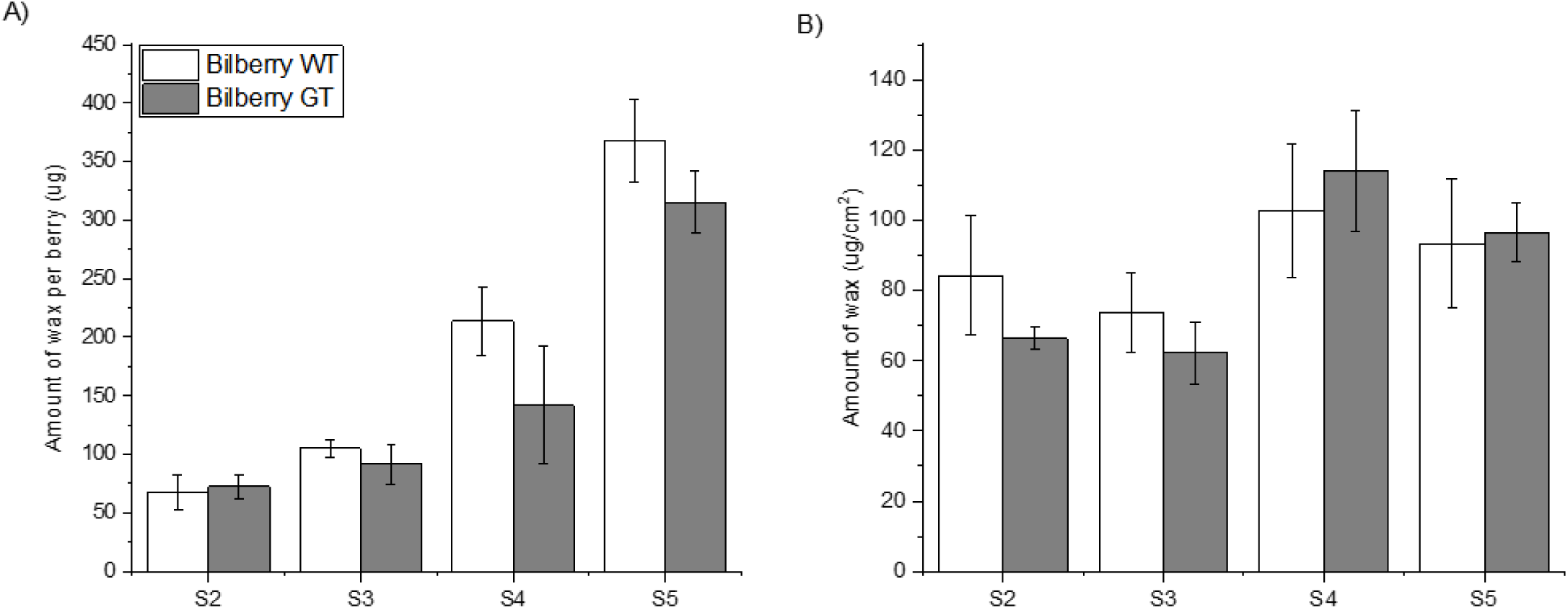
A) Amount of cuticular wax per berry fruit during ripening stages in wild type (WT) and glossy type mutant (GT) bilberry fruits (B) Amount of cuticular wax (in μg /cm^2^) in wild type (WT) and glossy type mutant (GT) bilberry fruits.

### Composition of cuticular wax

GC-MS analysis showed that the cuticular wax of both WT and GT fruit were mainly composed of triterpenoids, fatty acids, primary alcohols, ketones, aldehydes, and alkanes (Fig. 3). Triterpenoids followed by fatty acids were found to be the dominant compounds in all studied developmental stages of both WT and GT fruit cuticular wax. Secondary alcohols and esters were not detected in cuticular wax of either WT or GT fruit.

**Fig. 3.**
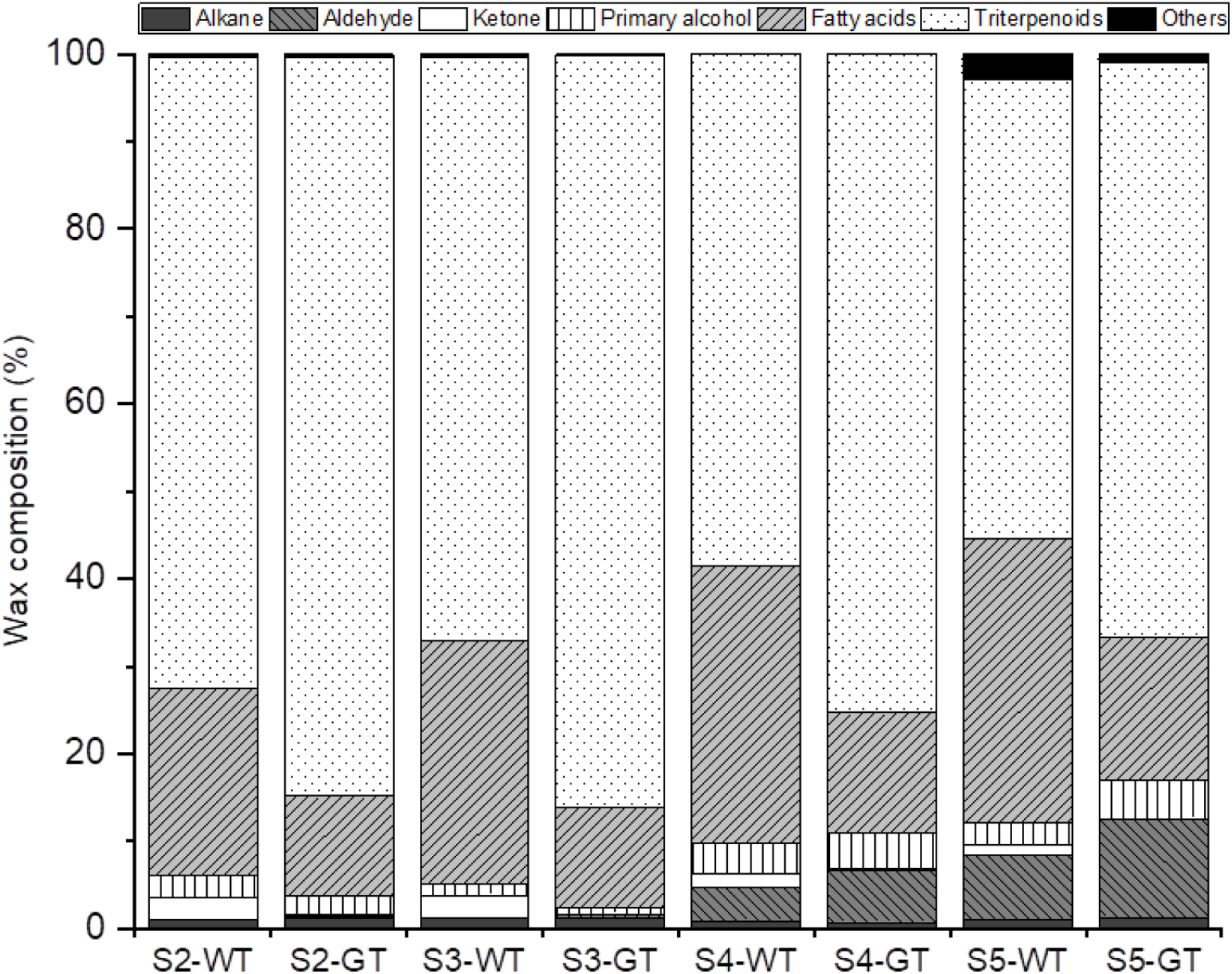
Proportion of chemical compound classes in wild type (WT) and glossy type mutant (GT) bilberry cuticular wax.

### Triterpenoids

The proportion of triterpenoids in cuticular wax showed differences through the course of bilberry fruit development as it was found to decrease from S2 to S5 (from 72.1% to 51.2%) in WT fruit (Fig. 3). Also in GT fruit cuticular wax, the proportion of triterpenoids was found to decrease during fruit development from S2 to S5 (from 84.5% to 65.0%). The triterpenoid proportion was higher in cuticular wax of GT fruit compared to WT fruit at all the studied stages of bilberry fruit development (Fig. 3). Relative triterpenoid proportion was found to be higher in GT fruit by 17% in S2, 29% in S3, 29% in S4 and 18% in S5 compared to WT fruits.

Generally, oleanoic acid was the predominant triterpenoid in cuticular wax of both WT and GT fruit during development (Table 1). Ursolic acid, β-amyrin, and α-amyrin were also found in all stages of WT and GT fruit cuticular wax. Lupeol was detected only in S3, S4 and S5 stage in both WT and GT berries. Levels of amyrins and lupeol were found to be highest in S4 stage. Esters of oleanane and ursane type triterpenoids were found specifically in S4 and S5 stage. Oleanoic acid was found in higher amounts in GT than WT fruit in S3, S4 and S5 stages. β-amyrin was present in higher amount in S2, S3 and S4 stage in GT than in WT fruits (Table 1).

**Table 1.**
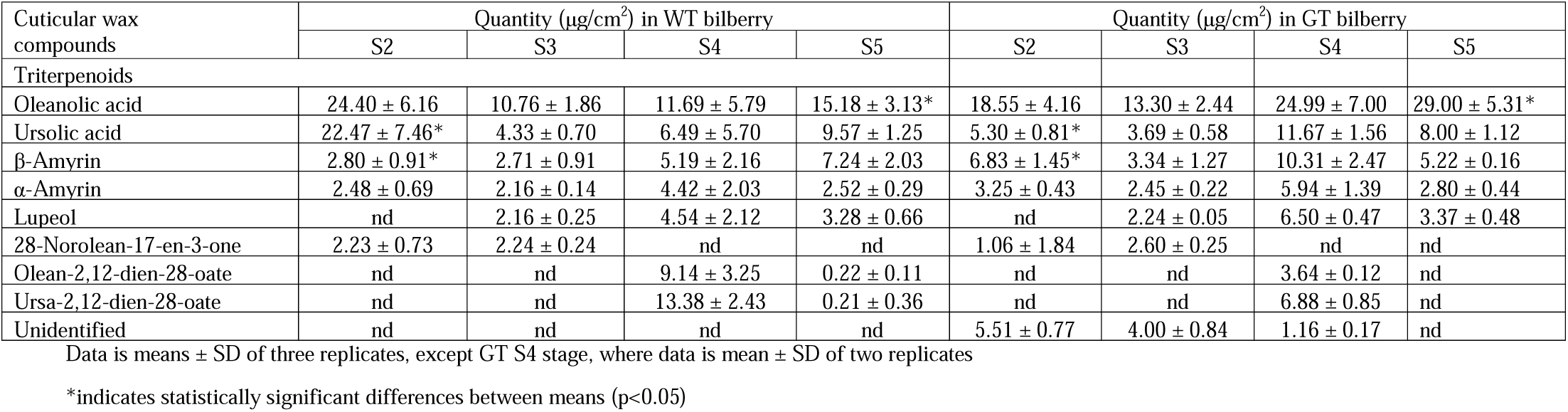
Quantities (μg/cm^2^) of triterpenoids during development of wild type bilberry (WT) and glossy type mutant (GT) fruits.

| Cuticular wax compounds | Quantity ($\mu\text{g}/\text{cm}^2$) in WT bilberry | | | | Quantity ($\mu\text{g}/\text{cm}^2$) in GT bilberry | | | |
| --- | --- | --- | --- | --- | --- | --- | --- | --- |
|  | S2 | S3 | S4 | S5 | S2 | S3 | S4 | S5 |
| Triterpenoids |  |  |  |  |  |  |  |  |
| Oleanolic acid | $24.40 \pm 6.16$ | $10.76 \pm 1.86$ | $11.69 \pm 5.79$ | $15.18 \pm 3.13^*$ | $18.55 \pm 4.16$ | $13.30 \pm 2.44$ | $24.99 \pm 7.00$ | $29.00 \pm 5.31^*$ |
| Ursolic acid | $22.47 \pm 7.46^*$ | $4.33 \pm 0.70$ | $6.49 \pm 5.70$ | $9.57 \pm 1.25$ | $5.30 \pm 0.81^*$ | $3.69 \pm 0.58$ | $11.67 \pm 1.56$ | $8.00 \pm 1.12$ |
| β-Amyrin | $2.80 \pm 0.91^*$ | $2.71 \pm 0.91$ | $5.19 \pm 2.16$ | $7.24 \pm 2.03$ | $6.83 \pm 1.45^*$ | $3.34 \pm 1.27$ | $10.31 \pm 2.47$ | $5.22 \pm 0.16$ |
| α-Amyrin | $2.48 \pm 0.69$ | $2.16 \pm 0.14$ | $4.42 \pm 2.03$ | $2.52 \pm 0.29$ | $3.25 \pm 0.43$ | $2.45 \pm 0.22$ | $5.94 \pm 1.39$ | $2.80 \pm 0.44$ |
| Lupeol | nd | $2.16 \pm 0.25$ | $4.54 \pm 2.12$ | $3.28 \pm 0.66$ | nd | $2.24 \pm 0.05$ | $6.50 \pm 0.47$ | $3.37 \pm 0.48$ |
| 28-Norolean-17-en-3-one | $2.23 \pm 0.73$ | $2.24 \pm 0.24$ | nd | nd | $1.06 \pm 1.84$ | $2.60 \pm 0.25$ | nd | nd |
| Olean-2,12-dien-28-oate | nd | nd | $9.14 \pm 3.25$ | $0.22 \pm 0.11$ | nd | nd | $3.64 \pm 0.12$ | nd |
| Ursa-2,12-dien-28-oate | nd | nd | $13.38 \pm 2.43$ | $0.21 \pm 0.36$ | nd | nd | $6.88 \pm 0.85$ | nd |
| Unidentified | nd | nd | nd | nd | $5.51 \pm 0.77$ | $4.00 \pm 0.84$ | $1.16 \pm 0.17$ | nd |

Data is means ± SD of three replicates, except GT S4 stage, where data is mean ± SD of two replicates
\*indicates statistically significant differences between means ( $p < 0.05$ )

### Aliphatic compounds

Generally, in both WT and GT fruits, the proportion of total aliphatic compounds increased during fruit development (Fig 3). A markedly lower proportion of total aliphatic compounds was observed in GT fruit relative to WT fruit in every developmental stage. This was mainly contributed by lower percentage of fatty acids in GT fruit compared to WT fruit (Fig 3). The proportion of fatty acids increased during both WT and GT fruit development. Montanic acid (C28) was the dominant fatty acid in both WT and GT fruits during S4 and S5 stages (Table 2).

**Table 2.** Quantities (μg/cm^2^) of very long chain aliphatic compounds during development of wild type bilberry (WT) and glossy type mutant (GT) fruits.

| Cuticular wax compounds | Quantity ($\mu\text{g}/\text{cm}^2$) in WT bilberry | | | | Quantity ($\mu\text{g}/\text{cm}^2$) in GT bilberry | | | |
| --- | --- | --- | --- | --- | --- | --- | --- | --- |
|  | S2 | S3 | S4 | S5 | S2 | S3 | S4 | S5 |
| *Fatty acids* |  |  |  |  |  |  |  |  |
| Oleic acid | $0.06 \pm 0.02$ | nd | $0.15 \pm 0.02$ | $0.06 \pm 0.01^*$ | $0.08 \pm 0.01$ | nd | $0.15 \pm 0.12$ | $0.13 \pm 0.03^*$ |
| Stearic acid | $0.39 \pm 0.01^*$ | $0.23 \pm 0.02^*$ | $0.57 \pm 0.08$ | $0.28 \pm 0.01^*$ | $0.17 \pm 0.01^*$ | $0.12 \pm 0.01^*$ | $0.25 \pm 0.18$ | $0.18 \pm 0.03^*$ |
| Nonadecanoic acid | $0.08 \pm 0.02$ | $0.05 \pm 0.04$ | nd | $0.05 \pm 0.04$ | nd | nd | $0.04 \pm 0.07$ | 0.00 |
| Arachidic acid | $8.34 \pm 0.40^*$ | $4.97 \pm 0.87^*$ | $6.40 \pm 0.29$ | $3.56 \pm 0.21^*$ | $1.07 \pm 0.20^*$ | $0.60 \pm 0.15^*$ | $1.47 \pm 1.25$ | $0.84 \pm 0.34^*$ |
| Behenic acid | $0.43 \pm 0.01^*$ | $0.18 \pm 0.04$ | $0.44 \pm 0.07$ | $0.29 \pm 0.04$ | $0.27 \pm 0.09^*$ | $0.19 \pm 0.05$ | $0.41 \pm 0.37$ | $0.36 \pm 0.06$ |
| Lignoceric acid | $0.54 \pm 0.09$ | $0.28 \pm 0.04$ | $0.62 \pm 0.11$ | $0.45 \pm 0.09$ | $0.47 \pm 0.13$ | $0.30 \pm 0.11$ | $0.45 \pm 0.34$ | $0.40 \pm 0.07$ |
| Hyenic acid | $0.11 \pm 0.04$ | $0.09 \pm 0.00$ | nd | $0.13 \pm 0.01$ | $0.11 \pm 0.00$ | $0.08 \pm 0.01$ | 0.00 | $0.12 \pm 0.01$ |
| Ceric acid | $1.60 \pm 0.47$ | $1.50 \pm 0.09^*$ | $3.40 \pm 0.55$ | $5.04 \pm 1.30$ | $1.10 \pm 0.23$ | $0.99 \pm 0.31^*$ | $1.88 \pm 1.33$ | $3.52 \pm 0.68$ |
| Carboceric acid | $0.07 \pm 0.02$ | $0.14 \pm 0.07$ | nd | $0.24 \pm 0.02^*$ | $0.08 \pm 0.00$ | $0.12 \pm 0.01$ | nd | $0.13 \pm 0.01^*$ |
| Montanic acid | $1.40 \pm 0.44$ | $1.99 \pm 0.23^*$ | $14.21 \pm 4.04$ | $10.37 \pm 2.00^*$ | $0.90 \pm 0.10^*$ | $0.94 \pm 0.01^*$ | $4.05 \pm 0.00$ | $5.03 \pm 0.01^*$ |
| Nonacosanoic acid | $0.07 \pm 0.02$ | $0.08 \pm 0.01$ | nd | $0.12 \pm 0.02$ | $0.04 \pm 0.07$ | nd | nd | $0.14 \pm 0.02$ |
| Melissic acid | $0.71 \pm 0.26$ | $0.60 \pm 0.10$ | $1.64 \pm 0.53$ | $1.67 \pm 0.94$ | $0.54 \pm 0.13$ | $0.51 \pm 0.26$ | $0.57 \pm 0.20$ | $0.92 \pm 0.32$ |
| *Ketones* |  |  |  |  |  |  |  |  |
| 2-Nonanone | $0.13 \pm 0.01^*$ | $0.05 \pm 0.03$ | nd | nd | $0.01 \pm 0.02^*$ | nd | nd | nd |
| 2-Undecanone | $0.04 \pm 0.01$ | $0.03 \pm 0.02$ | nd | nd | $0.04 \pm 0.00$ | nd | nd | nd |
| 2-Tridecanone | nd | nd | $0.12 \pm 0.11$ | $0.13 \pm 0.07$ | nd | nd | 0.04 | nd |
| 2-Nonadecanone | $0.05 \pm 0.01$ | $0.03 \pm 0.00$ | nd | nd | nd | nd | nd | nd |
| 2-Heneicosanone | $1.67 \pm 0.20$ | $0.92 \pm 0.24^*$ | $0.97 \pm 0.65$ | $0.64 \pm 0.05^*$ | $0.08 \pm 0.01$ | $0.05 \pm 0.00^*$ | $0.20 \pm 0.17$ | $0.04 \pm 0.01^*$ |
| 2-Docosanone | nd | nd | nd | nd | nd | nd | nd | nd |
| *Aldehydes* |  |  |  |  |  |  |  |  |
| Octadecanal | nd | nd | $0.03 \pm 0.03$ | $0.03 \pm 0.00$ | nd | nd | $0.01 \pm 0.01$ | $0.02 \pm 0.02$ |
| Tetracosanal | nd | nd | $0.05 \pm 0.04$ | $0.04 \pm 0.01^*$ | nd | nd | $0.06 \pm 0.03$ | $0.07 \pm 0.02^*$ |
| Pentacosanal | nd | nd | $0.07 \pm 0.02$ | $0.06 \pm 0.02$ | nd | nd | $0.05 \pm 0.04$ | $0.08 \pm 0.01$ |
| Hexacosanal | $0.04 \pm 0.03$ | $0.02 \pm 0.00$ | $1.03 \pm 0.07$ | $1.27 \pm 0.48^*$ | $0.03 \pm 0.00$ | $0.03 \pm 0.01$ | $1.25 \pm 1.13$ | $2.72 \pm 0.34^*$ |
| Heptacosanal | nd | nd | nd | $0.16 \pm 0.05$ | nd | $0.03 \pm 0.00$ | nd | $0.19 \pm 0.02$ |
| Octacosanal | $0.02 \pm 0.01$ | nd | $2.11 \pm 0.17$ | $3.35 \pm 0.72$ | nd | $0.04 \pm 0.00$ | $2.15 \pm 1.82$ | $4.65 \pm 0.45$ |
| Triacontanal | nd | nd | $0.25 \pm 0.24$ | $0.67 \pm 0.24$ | nd | nd | $0.28 \pm 0.11$ | $0.55 \pm 0.15$ |
| *Primary alcohols* |  |  |  |  |  |  |  |  |
| 1-Hexadecanol | $0.27 \pm 0.09$ | $0.25 \pm 0.07$ | $0.07 \pm 0.00$ | nd | $0.25 \pm 0.22$ | nd | nd | nd |
| 1-Octadecanol | $0.28 \pm 0.07^*$ | $0.38 \pm 0.01^*$ | $0.21 \pm 0.00$ | $0.29 \pm 0.00$ | $0.38 \pm 0.02^*$ | $0.29 \pm 0.03^*$ | nd | nd |
| 2-Octacosen-1-ol | nd | nd | nd | 0.27 ± 0.02 | nd | nd | nd | 0.38 ± 0.06 |
| 1-Eicosanol | nd | nd | 0.73 ± 0.10 | 0.23 ± 0.04 | nd | nd | 0.68 ± 0.06 | 0.33 ± 0.06 |
| 1-Docosanol | 0.16 ± 0.16 | nd | 0.74 ± 0.12 | 0.24 ± 0.04 | nd | nd | 0.67 ± 0.04 | 0.34 ± 0.06 |
| 1-Tricosanol | 0.15 ± 0.16 | nd | nd | nd | nd | nd | nd | nd |
| 1-Tetracosanol | 0.17 ± 0.16 | nd | 0.86 ± 0.20 | 0.30 ± 0.04 | nd | nd | 0.73 ± 0.05 | 0.40 ± 0.06 |
| 1-Pentacosanol | 0.05 ± 0.09 | nd | nd | 0.27 ± | nd | nd | nd | 0.42 ± 0.07 |
| 1-Hexacosanol | 0.05 ± 0.09 | nd | 0.83 ± 0.07 | 0.44 ± 0.06 | nd | nd | 0.90 ± 0.09 | 0.47 ± 0.08 |
| 1-Octacosanol | nd | nd | 0.29 ± 0.50 | nd | nd | nd | 0.82 ± 0.06 | 0.50 ± 0.07 |
| 2-Nonacosen-1-ol | nd | nd | nd | 0.26 ± 0.02 | nd | nd | nd | 0.38 ± 0.06 |
| *Alkanes* |  |  |  |  |  |  |  |  |
| Tetracosane | nd | 0.01 ± 0.02 | 0.25 ± 0.01 | 0.20 ± 0.04 | nd | 0.04 ± 0.00 | 0.23 ± 0.00 | 0.24 ± 0.09 |
| Pentacosane | 0.13 ± 0.04 | 0.09 ± 0.01 | 0.11 ± 0.01 | 0.04 ± 0.00 | 0.11 ± 0.00 | 0.09 ± 0.00 | 0.10 ± 0.02 | 0.09 ± 0.06 |
| Hexacosane | 0.04 ± 0.01 | 0.04 ± 0.00 | 0.51 ± 0.07 | 0.33 ± 0.24 | 0.06 ± 0.01 | 0.04 ± 0.00 | 0.41 ± 0.09 | 0.33 ± 0.12 |
| Heptacosane | 0.29 ± 0.09 | 0.18 ± 0.01 | nd | 0.03 ± 0.02 | 0.21 ± 0.05 | 0.16 ± 0.04 | nd | 0.06 ± 0.06 |
| Octacosane | 0.05 ± 0.01 | nd | nd | 0.11 ± 0.10 | nd | nd | nd | 0.09 ± 0.16 |
| Nonacosane | 0.22 ± 0.03 | 0.13 ± 0.04 | nd | nd | 0.18 ± 0.02 | 0.09 ± 0.00 | nd | 0.06 ± 0.05 |
| Hentriacontane | 0.09 ± 0.02 | 0.05 ± 0.02 | nd | nd | 0.08 ± 0.01 | 0.07 ± 0.02 | nd | 0.09 ± 0.04 |
| Total | 0.81 | 0.50 | 0.87 | 0.71 | 0.63 | 0.48 | 0.73 | 0.96 |
| *Phenolic acids* |  |  |  |  |  |  |  |  |
| Benzoic acid | 0.12 ± 0.04 | 0.10 ± 0.02 | nd | nd | 0.12 ± 0.03 | 0.09 ± 0.09 | nd | nd |
| p-coumaric acid | 0.05 ± 0.00 * | 0.01 ± 0.01 | nd | nd | 0.03 ± 0.00* | nd | nd | nd |

Data is means ± SD of three replicates, except GT S4 stage, where data is mean ± SD of two replicates
\*indicates statistically significant differences between means (p<0.05)

The proportion of ketones showed significant decrease in cuticular wax of GT fruit compared to WT fruit (Fig 3). The relative proportion decreased by 8 fold (S2), 19 fold (S3), 6 fold (S4) and 22 fold (S5) in GT than WT fruit. The proportion of ketones decreased slightly during WT fruit development. 2-heneicosanone (C21) was the dominant ketone found in both WT and GT fruit in all developmental stages but the amount was significantly higher in WT compared to GT fruit (Table 2).

Aldehydes were detected in high proportions only in S4 and S5 stages in both WT and GT fruit cuticular wax (Fig 3). Higher relative proportions of aldehydes were detected in GT compared to WT fruit by 53% in S4 and by 50% in S5 stage of fruit ripening. Octacosanal was the dominant aldehyde in both WT and GT fruits, followed by hexacosanal and triacontanal (Table 2).

Primary alcohols and alkanes showed a variable trend during development in both WT and GT fruits (Fig 3). A lower relative proportion of primary alcohols in GT relative to WT was observed with a decrease in S2 by 18% and S3 by 63%, followed by an increase in 11% in S4 and 74% in S5. Aromatic acids (phenolic acids) were found only in S2 and S3 developmental stages in both WT and GT fruit (Table 2).

### Identification and expression of cuticular wax biosynthetic genes

In the published bilberry transcriptome database (Nguyen *et al*., 2018), we were able to identify 335 unigenes encoding enzymes predicted to be involved in wax biosynthetic pathway, including fatty acid synthesis, fatty acid elongation, wax compound biosynthesis, wax transportation, and regulation of wax biosynthesis (Table S2). In the triterpenoid biosynthetic pathway, we identified 21 unigenes encoding two OSCs, namely BAS and LUS (Table S2). Sixteen unigenes were selected for gene expression analysis based on high sequence similarity with Arabidopsis and some fruit bearing species (Table S3).

The qRT-PCR results in pulp and skin of WT and GT fruits during development are shown in Fig. 4. Overall, the genes showed differential expression patterns during bilberry fruit development. Notably, the *CER26-like, FAR2, CER3-like, LTP, MIXTA*, and *BAS* genes were expressed at higher levels in the skin of both WT and GT fruits (Fig. 4).

**Fig. 4.**
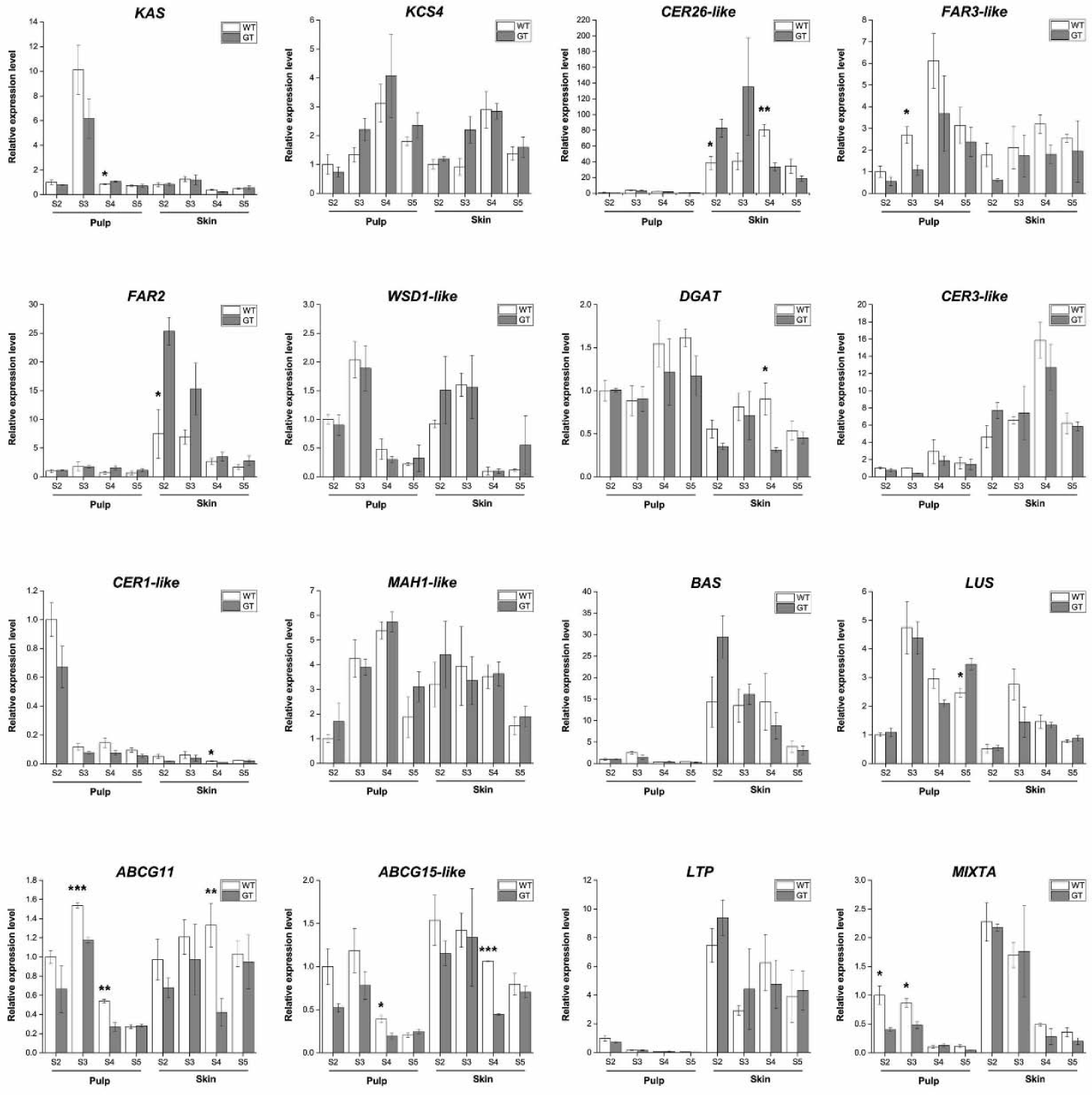
Gene expression of wax related genes in wild type bilberry (WT) and glossy type mutant (GT) were studied both in fruit pulp and skin during fruit development. S2, small green fruits; S3, large green fruits; S4, ripening red fruits; S5, fully ripe blue fruits. Error bars represent standard error of three biological replicates. The asterisks denote statistically significant differences between WT and GT (*: p≤0.05; **: p≤0.01; ***: p≤0.001).

In fatty acid biosynthetic pathway, bilberry unigene encoding KAS showed highest expression in pulp of both WT and GT fruits at developmental stage S3. In fatty acid elongation stage, *KCS4* transcript level was upregulated at the onset of ripening (S4) in both WT and GT fruits. Another elongation gene, *CER26-like* was predominantly expressed in the berry skin in both WT and GT fruits. Considering the differences in the gene expression of wax related genes in WT and GT bilberry fruit, we observed that the expression level of *CER26-like* was high at early stages in GT fruits in contrast to WT fruits which showed upregulation at the onset of ripening at S4 stage

In the alcohol-forming pathway, we identified a unigene annotated as *FAR3-like* in bilberry which was not found to be differentially expressed through all ripening stages between pulp and skin. However, *FAR2* exhibited skin-specific expression. The expression of *FAR2* gene was highest at development stages S2 and S3 and dramatically dropped thereafter in both WT and GT fruits. *FAR2* exhibited higher transcript abundance in GT than WT fruits. Two candidate genes encoding WSD1/DGAT showed no difference between pulp and skin in most of the developmental stages.

In the alkane-forming pathway, *CER3-like* was markedly up-regulated at the onset of ripening (S4) in both WT and GT fruits. In contrast, *CER1* did not differ in transcript levels in pulp and skin of WT and GT in the developmental stages except S4. *MAH1*, which has been related with the formation of the secondary alcohols, did not show differential expression between berry pulp and skin in developmental stages except S4.

In the triterpenoid biosynthetic pathway, *BAS* exhibited skin-specific expression in both WT and GT fruit. The expression pattern of *BAS* was high at early development stage S2, and was then gradually down-regulated throughout the ripening in GT fruit. The expression of *BAS* was also down-regulated at the fully ripe stage S5 in WT fruit. *LUS* gene showed higher expression in pulp with high expression at development stage S3.

Among the genes involved in the transportation of wax components, two *ABCG* genes, *ABCG11* and *ABCG15-like* were expressed higher levels in skin compared to pulp especially at ripening stages S4 and S5. *ABCG11* and *ABCG15-like* genes were down-regulated at the onset of ripening stage in GT skin and pulp compared to WT. Expression of *LTP* was found peaking at early development stage S2. The expression level of *LTP* gene was slightly higher in GT than WT bilberry. From the bilberry transcriptome database, we identified a unigene encoding MIXTA, a MYB transcription factor related to regulation of cuticle formation, which was up-regulated at early developmental stages S2 and S3.

*MIXTA* showed slightly higher expression level in WT than GT fruits in skin.

## Discussion

### WT and GT bilberry fruits both show accumulation of cuticular wax

Glossy, black bilberry mutant fruits have generally been considered to be waxless (Colak *et al*., 2017) although no scientific studies concerning the analysis of cuticular wax load has been reported previously. In orange, glaucous fruits have been demonstrated to contain higher cuticular wax load (Liu *et al*., 2012). However, in the present study, we found that both WT and GT bilberry fruits showed high and comparable accumulation of cuticular wax. Our results support the view that visual phenotype of plant cuticle is not correlated with the wax load (Adamski *et al*., 2013).

Based on our results, changes in wax biosynthesis and accumulation takes place during bilberry fruit development. Wax amount per berry increased during the fruit development of both WT and GT fruits indicating constant wax biosynthesis. Wax load per surface area remained somewhat constant due to growth of berry size although there were slight changes that can be attributed to the changes in the surface area compared to wax deposition rate. In other fruits, variable trends in wax load during fruit development have been reported (Trivedi *et al*., 2019b). Increase in wax load throughout the fruit development has also been reported in blueberry (Chu *et al*., 2018), apple (Ju and Bramlage, 2001), pear (Li *et al*., 2014) and orange fruits (Liu *et al*., 2012) whereas in grape, wax load increases until veraison followed by decrease in final ripening stage (Pensec *et al*., 2014).

### Glossy phenotype is attributed to changes in chemical composition affecting wax morphology

It has been previously reported that mutations in wax biosynthesis causing glossy surface in Arabidopsis leaf and stem show reduced density of wax crystals and sometimes also alterations in the crystal shape and size (Jenks *et al*., 1996). Similar results has been obtained in studies on surfaces of glossy fruits of orange (Liu *et al*., 2012; 2015) and cucumber (Wang *et al*., 2015a,b). Our study also demonstrated a decrease in the density of epicuticular wax crystal structures in GT fruit compared to WT fruit. While a dense cover of platelets along with rod-like structures were detected in S3, S4 and S5 stages in WT fruit, the surface of GT fruit was devoid of rod-like structures and dominated by membranous platelets. Our data suggest that the difference in appearance between WT and GT fruit of bilberry is based on the difference in epicuticular wax morphology that is due to differential chemical composition between WT and GT fruit.

Previously, Markstädter *et al*. (2000) correlated the glaucous phenotype stems of *Macaranga* species to higher triterpenoid content. In contrast, our study showed higher proportion of triterpenoids in glossy fruits compared to glaucous WT fruits. Since triterpenoids generally occur in intracuticular layer of wax (Jetter and Schaffer, 2001), they may not have a significant role in epicuticular wax crystal formation. Instead, epicuticular wax crystalloids are known to be dominated by aliphatic compounds. Previous studies have also attributed glaucousness to the presence of β-diketones in wheat flag leaf sheath (Zhang *et al*., 2013), however, in our study β-diketones were not found. Instead, among aliphatic compounds we observed the most prominent difference between WT and GT fruits in proportion of ketones. The result implies that glossy appearance in GT bilberry fruits could be due to the high reduction in amount of ketones. In supporting this hypothesis, our previous study showed that glaucous appearing bilberry (rod-like epicuticular morphology) and bog bilberry (coiled rodlet morphology) contain ketones while glossy appearing lingonberry and crowberry are devoid of ketones as well as rod-like structures (Trivedi *et al*., 2019a). Ketones have earlier been reported to be responsible for the formation of transversely rigid rodlets (Meusel *et al*., 1999). Also, cuticular waxes including ketones have been reported to form different types of rodlets in different plant species (Ensikat *et al*., 2006).

### Chemical composition of cuticular wax changes during bilberry fruit development

The chemical composition of ripe WT bilberry fruit cuticular wax corroborates with our previous study (Trivedi *et al.*, 2019a). However, the wax composition showed changes during the course of bilberry fruit development with the proportion of major compound classes generally varying similarly in both WT and GT fruits. A decrease in the proportion of triterpenoids and an increase in proportion of total aliphatic compounds was detected during bilberry fruit development. The decrease in the proportion of triterpenoids during fruit development has also been reported in grape (Pensec *et al*., 2014) and sweet cherry (Peschel *et al*., 2007). In accordance to our study, a recent study in bilberry reported lowest percentage of triterpenoids in cuticular wax of young fruits with increase during fruit development (Dashbaldan *et al*., 2019). However, in blueberry fruits the proportion of triterpenoids increased through developmental stages (Chu *et al*., 2018) indicating differences in wax biosynthesis even between closely related species. During bilberry fruit development, the presence of aldehydes during the later stages of berry development (S4 and S5) indicates that these are the key stages for biosynthesis of aldehydes in bilberry fruit cuticular wax. In wax biosynthetic pathway (Fig. S1), secondary alcohols are precursors for ketones, however, secondary alcohols were not observed in bilberry cuticular wax. The formation of ketones without the formation of secondary alcohols remains elusive. This might suggest that secondary alcohols are converted directly to ketones in bilberry or that ketones are biosynthesized via a different pathway in bilberry compared to Arabidopsis but needs further studies.

### Role of wax biosynthetic genes in bilberry fruit cuticular wax formation

The genes proposed to be involved in wax biosynthesis in bilberry showed differential expression profiles through the course of fruit development with markedly different expression of some genes in skin compared to pulp indicating their attendance in wax biosynthesis into cuticle.

Our results demonstrated uniform gene expression of *KAS* gene in the studied bilberry fruit tissues (skin and pulp) attributed to the broad role of KAS in synthesis of *de novo* fatty acid precursors, which can be partitioned to various pathways, such as suberin and cutin (Samuels *et al*., 2008). *KAS* expression profile is in line with our observation that the fatty acids proportion increases through the course of development gradually. The highest amounts of fatty acid precursors detected in S3 stage is most likely followed by further distribution of precursors to different wax biosynthesis pathways. The high upregulation in *KAS* gene expression at S3 in pulp in both WT and GT berries may indicate high fatty acid biosynthesis in bilberry seeds for synthesis of seed oils at S3 stage. It has been shown that bilberry seed oil has high content of PUFAs (C18) and vitamin E (Yang *et al*., 2011; Gustinelli *et al*., 2018).

For the fatty acid elongation, 21 *KCS* genes have been identified in Arabidopsis of which several genes were proposed to have roles in determining specific chain length of VLCFAs in different organs (Tresch *et al*., 2012). The transcript level of unigene for bilberry KCS4 was up-regulated at the onset of ripening (S4) in WT and GT fruits whereas bilberry *CER26-like* gene had highest expression already at S3 stage in GT fruit. *CER26-like* gene has been characterized for the elongation of specific chain length longer than C_28_ in leaves and stem of Arabidopsis (Pascal *et al*., 2013). The skin-specific expression of *CER26-like* gene suggests that it may play an important role in biosynthesis of very long chain fatty acids (VLCFAs) and its derivatives in bilberry. The differential expression of *CER26-like* genes between WT and GT fruit skin suggests that this gene might be responsible for differential accumulation of very long chain aliphatic compounds.

We observed the skin specific expression of *FAR2* (Fig. 4), a homolog of *AtFAR2* that produces primary alcohols incorporated into sporopollenin of the pollen exine layer (Chai *et al*., 2018). This suggests the role of *FAR2* gene in alcohol forming pathway in bilberry fruit.

In Arabidopsis, mutation of *CER3* gene led to a decrease in the amount of aldehydes, alkanes and their derivatives (Rowland *et al*., 2007). In bilberry, the accumulation trend of aldehydes in cuticular wax corroborated with the gene expression trend of *CER3-like* gene, both increasing at late ripening stages. This is in accordance with the expression pattern of *CER3* during fruit ripening of sweet cherry, mango, and orange (Alkio *et al*., 2012; Wang *et al*., 2016; Tafolla-Arellano *et al*., 2017). Therefore, we hypothesize that in bilberry *CER3* gene is involved in biosynthesis of aldehydes.

The intracellular transport of wax compounds from ER to plasma membrane is proposed to occur either by trafficking through Golgi system (McFarlane *et al*., 2014), or by oil bodies in the cytoplasm (Li *et al*., 2016). It is well established that ABCG transporters are required for wax transport across the plasma membrane (McFarlane *et al*., 2010). Lipid transfer proteins are also responsible for transporting lipid compounds in the cell wall. In Arabidopsis, *ABCG11* and *ABCG12* have been identified and characterized for function in wax deposition in stem (McFarlane *et al*., 2010). In the present study, we found higher expression of *ABCG15-like* in fruit skin suggesting that this gene may play a role in the wax transport in bilberry cuticle. Similarly, skin-specific expression of *LTP* gene in bilberry suggests its role in transportation of wax compounds in the fruit cuticle.

In fleshy fruits, some regulatory genes of cuticular wax biosynthesis have been identified and characterized e.g. *MdSHN3* in apple (Lashbrooke *et al*., 2015b), tomato *SlSHINE3* and *SlMIXTA*, a MYB regulator downstream to *SlSHINE3* (Shi *et al*., 2013; Lashbrooke *et al*., 2015a). These positive regulators have been proposed to affect cuticle formation and epidermal cell differentiation (Oshima *et al*., 2013; Lashbrooke *et al*., 2015a). *SlMIXTA* has been shown to be down-regulated during tomato fruit ripening (Lashbrooke *et al*., 2015a) similar to the qRT-PCR results of this gene in bilberry fruits. Therefore, our results suggest that the MIXTA plays a role in the cuticle of bilberry fruits at early developmental stages.

The cuticular wax pathway has been characterized in plants, however the biosynthesis and transport of triterpenoids in cuticular wax is a topic less explored. We observed skin specific expression of *BAS* in bilberry fruit skin, similar to two *OSC* genes in tomato, *SlTTS1* and *SlTTS2*, which were expressed exclusively in the epidermis and produced triterpenoids for the fruit cuticular wax (Wang *et al*., 2011). The high expression of *BAS* in early stage of development is in line with the high expression of triterpenoids generally in early stages of development.

## Conclusions

Based on our results, bilberry GT fruits have cuticular wax load comparable to WT bilberry fruit. However, the chemical composition and morphology of cuticular wax along with gene expression for wax biosynthetic genes varied between GT fruit and WT fruit. GT fruit had higher content of triterpenoids accompanied by lower content of fatty acids, ketones compared to WT fruit. Significant reduction of ketones was accompanied by the loss of rod-like structures in GT fruit cuticular wax suggest a correlation between glaucousness and ketones in bilberry fruit cuticular wax. The skin specific expression of *CER26-like, FAR2, CER3-like, LTP, MIXTA-* and *BAS* underlines the role of these genes in wax biosynthesis in bilberry.

## Abbreviations

BAS: β-amyrin synthase
KAS: β-ketoacyl-ACP synthase
KCS: β–ketoacyl-CoA-synthase
FAR: Fatty acyl-CoA reductase
GT: Glossy type mutant
LTP: Lipid transfer protein
LUS: Lupeol synthase
MAH1: Mid-chain alkane hydrolase
DGAT: Diacylglycerol acyltransferase
OSCs: Oxidosqualene cyclase enzymes
SEM: Scanning electron microscopy
VLCFAs: Very long chain fatty acids
WSD1: Wax synthase
WT: Wild type

## Supplementary data

Table S1. Primers used for qRT-PCR analysis.

Table S2. Number of unigenes involved in the cuticular wax biosynthesis of bilberry.

Table S3. Characterization of wax-related genes in bilberry.

Fig. S1. Schematic presentation of cuticular wax biosynthetic pathway. PM: plasma membrane, CW: cell wall.

## Acknowledgments

This work was financially supported by I4 future doctoral program, hosted at the University of Oulu: Novel Imaging and Characterization Methods in Bio, Medical, and Environmental Research and Technology Innovations, which is the European Union’s Horizon 2020 Research and Innovation Programme under the Marie Sklodowska-Curie action co-funded by international, interdisciplinary and inter-sectoral doctoral programme (grant number 713606 to PT). The research was also funded by InterregNord (Natural Wax of Arctic Berries as Our Treasure – WAX project (number 20201089 to University of Oulu and grant IR16-020 and grant RMF16-026 to Troms Fylkeskommune and NIBIO).

